# Modulation of visually induced self-motion illusions by α transcranial electric stimulation over the superior parietal cortex

**DOI:** 10.1101/2023.05.22.540983

**Authors:** Sylvain Harquel, Corinne Cian, Laurent Torlay, Emilie Cousin, Pierre-Alain Barraud, Thierry Bougerol, Michel Guerraz

**Author notes:** corresponding author Sylvain Harquel, UPHUMMEL, Defitech Chair of Clinical Neuroengineering, Neuro-X Institute (INX) and Brain Mind Institute, École Polytechnique Fédérale de Lausanne (EPFL), 9 Chemin des Mines, 1202 Geneva, Switzerland.

## Abstract

Growing popularity of virtual reality systems has led to a renewed interest in understanding the neurophysiological correlates of the illusion of self-motion (vection), a phenomenon that can be both intentionally induced or avoided in such systems, depending on the application. Recent research has highlighted the modulation of α power oscillations over the superior parietal cortex during vection, suggesting the occurrence of inhibitory mechanisms in the sensorimotor and vestibular functional networks to resolve the inherent visuo-vestibular conflict. The present study aims to further explore this relationship and investigate whether neuromodulating these waves could causally affect the quality of vection. In a crossover design, 22 healthy volunteers received 13 minutes of high-amplitude, focused α-tACS over the superior parietal cortex while experiencing visually induced vection triggered by optokinetic stimulation. The tACS was tuned to each participant’s individual α peak frequency, with θ-tACS and sham stimulation serving as controls. Overall, participants experienced better quality vection during α-tACS compared to control θ-tACS and sham stimulations, as quantified by the intensity of vection. The observed neuromodulation supports a causal relationship between parietal α oscillations and visually induced self-motion illusions, with their entrainment triggering over-inhibition of the conflict within the sensorimotor and vestibular functional networks. These results confirm the potential of non-invasive brain stimulation for modulating visuo-vestibular conflicts, which could help to enhance the sense of presence in virtual reality environments.

## Introduction

The optic flow caused by relative motion between an observer and the visual environment can induce an illusory sensation of passive self-motion, referred to as vection (Dichgans & Brandt, 1978). When a stationary observer views a large scene moving uniformly in one direction, he/she soon feels himself moving in the opposite direction. The scene itself may appear to slow down or even stop moving (Dichgans & Brandt, 1978; Guerraz & Bronstein, 2008; Howard, 1982). It has been suggested that the occurrence of vection is related to the *a priori* assumption of a stable environment, making it more likely that motion cues are attributed to self-motion (Dichgans & Brandt, 1978; Rock, 1990). However, since vection is induced entirely by visual motion, a sensory conflict occurs between visual information and the graviceptive one that signals the absence of self-motion (particularly vestibular information; Cheung et al., 1989). It has been further suggested that an optimal multisensory integration involves higher weighting of more reliable signals (Ernst & Banks, 2002). Thus, the sensory conflict must be resolved by down-weighting vestibular cues and up-weighting visual cues. Accordingly, positron emission tomography and functional Magnetic Resonance Imaging (fMRI) studies in humans (Brandt, 1998; Deutschländer et al., 2002, 2004; Dieterich et al., 2003; Kleinschmidt et al., 2002) have shown that vection-generating continuous optokinetic stimulation not only activates the visual cortex but also concomitantly deactivates a parieto-insular region referred to as the “vestibular cortex”, which is part of a wider cortical and subcortical vestibular network (Dieterich & Brandt, 2018; Guldin & Grüsser, 1998; Lopez et al., 2012). Deactivation of vestibular areas may reflect the latter’s inhibition by the visual system (Brandt, 1998), in order to resolve conflicting (incongruent) inputs (Brandt & Dieterich, 1999; Brandt et al., 2002). However, the level of deactivation does not result only from the continuous optokinetic stimulation *per se*, but also from its perceptual interpretation. Indeed, Kleinschmidt *et al*. (2002) showed that the percept of self-motion as compared to the percept of object motion, also modulated neural activity with 1) a lower level of neural activity in the visual areas (such as in V5/MT, V4, V3/V3a, V1) and 2) an even lower activity in the vestibular areas (Kleinschmidt et al., 2002). More recently, the literature has focused on the electrophysiological mechanisms supporting this phenomenon, and on its exact timing relative to the occurrence of vection.

Electroencephalography (EEG) studies have shown that vestibular processing and self-motion estimates are related to α power modulation. On one hand, vestibular stimulation induced by passive whole-body motion is associated with the suppression of α power (8-13 Hz) mainly over the left and right temporoparietal regions (Gale et al., 2016; Gutteling & Medendorp, 2016). On the other hand, an increase in α activity over the superior parietal cortex has been found during visually induced vection (Harquel et al., 2019; McAssey et al., 2020; Palmisano et al., 2016) that correlated with the strength of the vection experience (Harquel et al., 2019; Palmisano et al., 2016). Furthermore, the precise timing of such α power modulations was characterized by a transient and short-lasting decrease around vection onset followed by a sustained increase (i.e. above the baseline measured in the absence of self-motion perception) (Harquel et al., 2019; McAssey et al., 2020). Indeed, the elevated α power was maintained throughout the perception of self-motion and only returned to the baseline level at the offset of vection (Harquel et al., 2019). Overall, it has been suggested that α oscillations reflect functional inhibition, irrelevant information being filtered out by increasing the α power in the areas processing this information (for a review, see Van Diepen et al., 2019). In terms of vection, the modulation of α activity may reflect the level of inhibition in the vestibular network as previously reported in fMRI studies, blood oxygen level-dependent signal being negatively correlated with α activity (Feige et al., 2005; Goldman et al., 2002; Laufs et al., 2003; Scheeringa et al., 2009). This inhibitory process is needed to reduce potential interference from conflicting vestibular inputs that would otherwise reduce vection experiences (Dichgans & Brandt, 1978). This is in agreement with the increased subjective intensity of vection reported by labyrinthine defective subjects in whom there is no more possible interference from conflicting vestibular inputs (Johnson et al., 1999). This hypothesis is also corroborated by the differences in α power modulation as a function of the visual-vestibular conflict, as observed for visual motion rotating about the line of sight in standing or supine healthy observers (Harquel et al., 2019). Since visual roll motion induces a persistent visual-otolith conflict only when the observer is standing (for a review, see Tanahashi et al., 2012), the additional increase in α activity specifically observed in this position underlined the level of inhibition needed to reduce further interference from conflicting vestibular inputs (Harquel et al., 2019). Accordingly, α activity might reflect the degree of discrepancy between actual and expected vestibular activities.

This converging evidence suggests that modulation in α activity may be a good marker of the conflict between sensory inputs, and might even be predictive of the occurrence of self-motion illusion. However, the above-mentioned correlation studies are limited because they rely on correlational inference between the electrophysiologcial readouts (α power) and the illusion, and causal evidence about this link is currently lacking (Herrmann et al., 2016). In the present study, the putative causal nature of this relationship was determined by using non-invasive brain stimulation (Herrmann et al., 2013; Thut et al., 2017). Specifically, transcranial alternating current stimulation (tACS) technique has been proven to modulate ongoing oscillatory brain activity by applying an alternative electrical current on the scalp surface (Antal et al., 2008; Dowsett et al., 2020; Helfrich et al., 2014; Neuling et al., 2015; Ruhnau et al., 2016; Vossen et al., 2015; Witkowski et al., 2016). Here, we targeted the superior parietal cortex, as an entry point to the vestibular functional network, using high-amplitude α frequency tACS (α-tACS) (Khatoun et al., 2018; Schutter & Wischnewski, 2016) on healthy participants to establish whether the modulation of α oscillations causally impacts illusory self-motion perception. To this end, this study was performed in a randomized, controlled, crossover design to assess the specific effects of α-tACS on roll vection in real time. α-tACS, θ-tACS (4-7 Hz) and sham stimulations were blindly delivered to the participant within a single experimental session. The α-tACS parameters were based on each participant’s individual α peak frequency (IAF). Self-motion illusions were evoked by the use of an optokinetic roll stimulation, known to induce alternating periods during which participants reported either a sensation of self-motion or a perception of themselves as static. This situation of bistable perception allows us to investigate the intensity of the subjective experience but also its occurrence and duration. We hypothesized that the modulation of α oscillations by non-invasive electrical stimulation would specifically modulate vection quality, assessed by vection duration and intensity, thus attesting to a causal relationship between α oscillations modulation in the vestibular functional network and visually induced self-motion illusions.

## Materials & Methods

### Participants

The sample size was computed using G-power 3.1.9.7 (Faul et al., 2009). Sample size was calculated in order to achieve 90% of power using an ANOVA (within subject design). Parameters for the *a priori* power analysis were: effect size set to 0.5 (corresponding to a partial η_p_^2^ of 0.2, very large effect), α set to 0.05, one group, three measurements, and the default parameters for the correlation between measurements (0.5) and nonsphericity correction (0.75). The effect’s size prior was evaluated to a very large effect of 0.2 (Cohen, 1988; Sawilowsky, 2009), based on two previous studies comparing vection duration and intensity in different experimental conditions (Harquel et al., 2019; McAssey et al., 2020), which are the most comparable to ours in terms of protocol design, especially regarding the vection induction procedure and measured outcomes. McAssey *et al*. (2020) found huge effect sizes when comparing vection duration and subjective intensity (Cohen’s d ≈ 2.8 and 2, respectively) during coherent versus incoherent motions. In our previous study (Harquel et al., 2019), we found large effect sizes (Cohen’s d ≈ 1.1) when using subjective scales in comparing vection intensity across two experimental conditions. However, it must be noted that the huge effects reported by McAssey *et al*. (2020) came from the strong difference between visual coherent and incoherent stimuli that were specifically designed to either trigger or prevent vection. Such effect sizes are overrated in the present context of a *neuromodulation* effect. The sample size required to achieve such statistical power was 13. The final sample size was nearly doubled to 24, in order to allow for the proper counterbalancing of our experimental conditions and to account for putative drop-outs.

Twenty-four healthy young adults (10 women, 14 men; age: 24 ± 3.76 y.o.; age range: 19 to 33 y.o.) participated in the study. They were all right-handed and had normal or corrected-to-normal vision. Handedness was ascertained by the use of the 10-item Edinburgh Handedness Inventory (EHI; Oldfield, 1971). Only participants showing a score above 50% of right handedness were considered for this study; the 50% cut-off was based upon work by Dragovic (2004). Individuals with vestibular or central nervous system disorders or any other contraindication to tACS and MRI were not included. Two participants had to be discarded from further analysis, due to a vasovagal episode during the familiarization phase or to a misunderstanding of the behavioral task, leading to a final number of 22 participants (8 women, 14 men; age: 24.23, 3.78 y.o.; age range: 20 to 33 y.o.). The study has been approved by the local investigational review board (Grenoble University Hospital, Grenoble, France; reference: ID RCB 2019-A03159-48). In accordance with the Declaration of Helsinki, all participants gave their informed written consent prior to their participation. The participants received financial compensation. MRI, tACS and EEG acquisitions were performed at IRMaGe MRI and neurophysiology facilities (Grenoble, France).

### Experimental design

tACS was administered using a randomized crossover design with three stimulation conditions: α-tACS, θ-tACS and sham stimulation, the last two serving as controls. Participants were informed that they would be equally stimulated during the three stimulation conditions, and that skin perception of this stimulation may vary from one condition to another. Before the stimulation sessions, the IAF, the stimulation discomfort during α-tACS applied with 1.5 mA/1 cm² current, and the presence or absence of phosphenes were evaluated. The IAF was then determined from spontaneous EEG activity with the eyes closed by identifying each individual peak frequency in the α-range (8–12Hz) over the parietal area using Fast Fourier Transforms (Corcoran et al., 2018; Klimesch et al., 1990). To evaluate any possible discomforting scalp sensation, participants were asked to report the presence of unpleasant symptoms like tingling, itching or burning. Next, the three tACS sessions (α-tACS, θ-tACS and sham) combined with optokinetic stimulation were administered in a counterbalanced order (Fig. 1).

**Figure 1.**
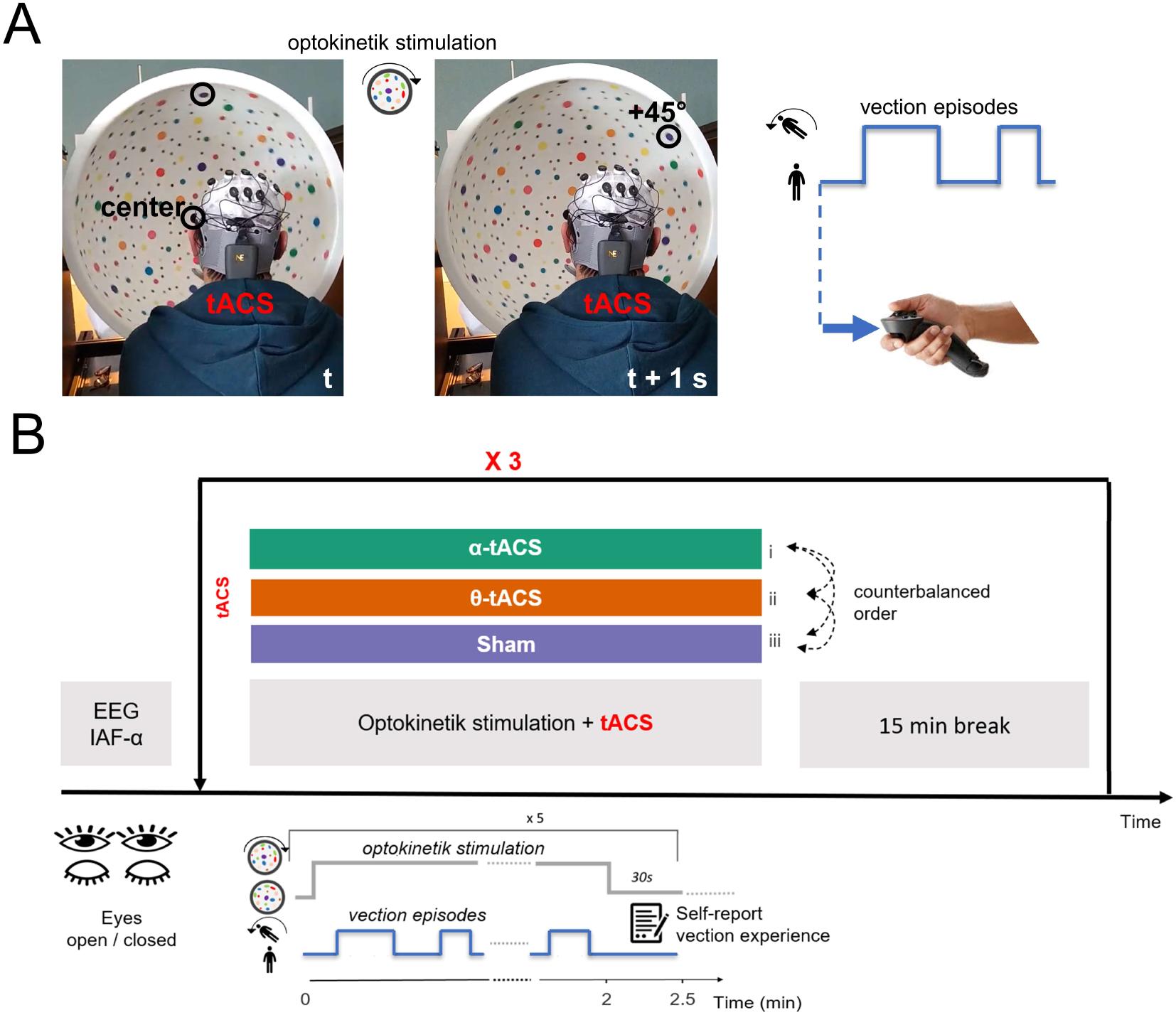
Experimental design and procedures. **A**: Time-lapse (1 s between the two snapshots) of the experimental apparatus showing the rotating dome used for optokinetic stimulation, and the tACS system (left, for details see Fig. 2). The participant was slightly offset to the right side, and the lighting was brighter for the needs of the photos. Example of behavioral responses: whenever the participants had the feeling of vection, they had to push a button with their right index finger and hold it until the end of the illusion (right). **B**: After preparatory EEG measurements (IAF evaluation), the experimental protocol consists of three optokinetic stimulation sequences with concurrent tACS stimulation. Three tACS stimulation conditions were tested in a counterbalanced order between participants, which are α-tACS and two control stimulation conditions: θ-tACS and sham.

Each session included five runs lasting 2 min with optokinetic stimulation. The first run was considered as a tACS familiarization run, while the subsequent four were considered as experimental runs. There was a 30 s period with a stationary dome after each run. The participants performed the perceptual task in an upright position (seated, with their head on a chin rest). The dome’s axis of rotation was aligned with their line of sight. The participants were instructed to (i) fixate the black spot in the center of the dome, (ii) avoid blinking, and (iii) not follow any moving dots with their eyes. Whenever the participants had the feeling of an illusory self-motion in roll (vection), they pushed a button with the index finger on a joystick held in their right hand and held it until the end of the illusion (see Fig. 1A). Since expectation about plausibility of real body motion may alter vection (D’Amour et al., 2021), participants were fully aware that self-motion perception was illusory. Earlier studies of circular vection established that a visual motion stimulus generates unstable perception with alternating periods of self-motion and no self-motion perception, and that participant can maintain fixation for several minutes (Harquel et al., 2019; Kleinschmidt et al., 2002; Seno et al., 2018; Thilo et al., 1999). After each run, they verbally reported the strength of their vection experience on a scale of 0 (“*no vection*”) to 10 (*“I felt I was really moving*”) (McAssey et al., 2020). The responses were recorded by the experimenter before the next run. The participants were familiarized with the perceptual task and the visual stimulus in a separate session prior to the stimulation sessions.

Each session ended with the participants’ reports on the intensity of unpleasant skin sensations during stimulation and any sensation of motion sickness. The unpleasant skin sensations were rated using a Likert scale ranging from 1 (“*no particular sensation*”) to 5 (“*very unpleasant to the point of wanting to stop the experience*”). To ensure that the participant did not suffer from motion sickness, the fast motion sickness scale (FMS; Keshavarz & Hecht, 2011) was administered. The FMS is a subjective rating scale of motion sickness ranging from 0 (“*no sickness*”) to 20 (“*severe sickness*”). Each stimulation session was followed by a 15 min break. Finally, a cerebral anatomic T1-weighted MRI was performed in order to obtain an individual estimation of the electric field strength (Kasten et al., 2019) using software modeling (Thielscher et al., 2015).

## Materials

### Optokinetic stimulation

The visual motion stimulation was performed using a fiberglass dome (diameter: 58 cm; see Fig. 1A and Harquel et al., 2019). The inside is painted white and is covered with a random pattern of dots of different diameters (from 0.5 cm to 2 cm) and colors (red, blue, orange, yellow, green, and purple). The dots account for around 40% of the total surface area. A black spot at the center of the dome provides a fixation point. The dome can be rotated around its center in a clockwise direction (roll plane) from the participant’s viewpoint by a 12 V electric motor (speed: 45 °/s). The apex of the dome was maintained at a distance of 26 cm from the viewer’s nasion. The visual field subtended a visual angle of 220°, that is, the entire visual field, so that no vertical or horizontal features of the experimental room can be seen. The experiment was carried out in an illuminated environment (2.05 cd/m^2^ for the white background of the dome), allowing the participant to see the color dots very clearly.

### Transcranial electrical stimulation

An integrated tES-EEG system (Starstim System, Neuroelectrics Barcelona SL, Spain) was used to deliver tACS, whose final montage is depicted in Fig. 2A. Stimulation was applied via small, round gel electrodes (Neuroelectrics Pistim Ag/AgCl, 3.14 cm^2^) positioned inside a non-conductive neoprene cap according to the international 10–10 EEG system (Jurcak et al., 2007). Each of the return and input electrodes was filled with a conductive gel keeping impedances below 10 kΩ. Using high-definition and high-amplitude tACS, our aim was to induce an electrical field centered on the superior parietal cortex, covering the bilateral superior parietal lobes (Brodmann area – BA 7) and the primary somatosensory cortex (S1, BA 5) (see Fig. 1C), where the α modulations during vection were previously observed (Harquel et al., 2019; McAssey et al., 2020). We paid special attention to maintain a maximum electric field onto the targets. This focality is important to spare neighboring cortical sites from undesired stimulation. Moreover, focal high-amplitude tACS montages are reported to avoid secondary effects such as phosphenes or skin irritations (Khatoun et al., 2018). In order to create such a montage on both hemispheres, we used SimNIBS software version 4.0 (Thielscher et al., 2015) tES modeling software with its brain template and tried numerous combinations of 2 x 3 electrodes including algorithmic optimization (Ruffini et al., 2014). The best montage in terms of focality and intensity was achieved by placing six gelled Ag/AgCl return electrodes with a contact area of π cm^2^ (NG PiStim, Neuroelectrics) at P1 /1.5 mA (0°), P2 / 1.5 mA (0°), C1 / 0.75 mA (180°), C2 / 0.75 mA (180°), C3 / 0.75 mA (180°) and C4 / 0.75 mA (180°) sites (10-10 EEG system, see Fig. 2A). This montage leads to significant electrical fields on the cortical targets peaking at 0.5 V/m, i.e. more than twice the minimum electric field of 0.2 V/m able, *in vitro*, to induce an effect (Reato et al., 2010).

**Figure 2.**
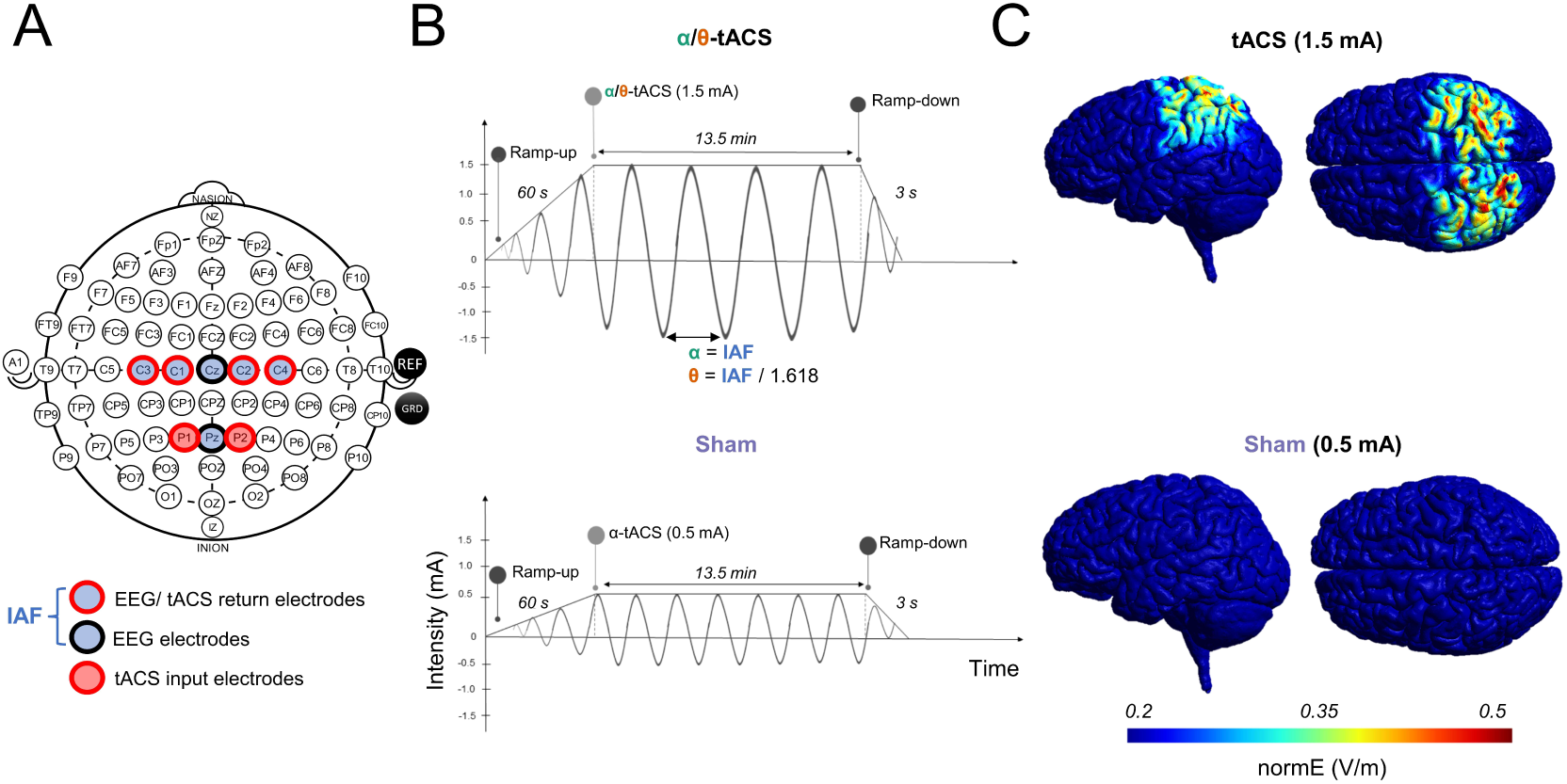
tACS/EEG electrode configuration and simulated electrical field distribution on a brain template. **A**: HD multi-electrode stimulation montage showing 2 input electrodes and 4 return electrodes placed over centroparietal scalp sites. Location of 6 EEG recording electrodes used for the IAF (C1, C2, C3 C4, Cz and Pz) according to the international 10–10 system is also depicted. **B:** Schematic representation of the tACS protocol. Electrical stimulation increases for a period of 60 seconds, is sustained for 1 minute, before the 12.5 min optokinetic stimulation protocol begins, and then decreases for 3 seconds. The maximum peak amplitude of tACS is set at 1.5 mA for active tACS condition (top row), and at 0.5 mA for sham (bottom row). **C:** Head model depicting the distribution of the E-field (2 inputs at 1.5mA, 4 returns diphase at 0.75mA) in the brain (1cm2 pi-electrodes) while stimulating with active tACS (top row) and sham (bottom row). The modeling of cortical current shows high bilateral focality of the electrical field in the superior parietal cortex (BA 5 & BA 7).

According to literature reports, the amplitude of α activity is enhanced when neural populations are stimulated at their resonance frequency (Merlet et al., 2013; Zaehle et al., 2010). Thus, α-tACS was applied at the participant’s IAF frequency. Stimulation duration (amount of on-time) in each active tACS and sham conditions lasted 14 min and 3 sec (Fig. 2B). Such amount of time does not exceed the limit values of maximum stimulation duration recommendations in literature and safety guide of Starstim neuroelectrics device (Antal et al., 2017; Nitsche et al., 2003). The applied current intensity was 1.5 mA / 1 cm² (3 mA peak-to-peak). The current was ramped up over the first 60 s and down over the last 3 s (Fig. 2B). For the θ-tACS stimulation, the same parameters were applied except the stimulation frequency, which was tuned relative to the participant’s IAF and set as: IAF / 1.618. Such a ratio theoretically prevents any correlation between the two stimulation frequencies (Pletzer et al., 2010). For the sham stimulation, α-tACS was applied at the participant’s IAF frequency with a small intensity (0.5 mA / 1 cm² - 1 mA peak-to-peak), designing to mimic skin effects without any efficient electrical field created in the cortex (which peaked at 0.16 V/m, i.e. 20% below the minimum efficiency threshold of 0.2 V/m, see Fig. 2C).

### Data analysis

#### Behavioral data

Three vection parameters were calculated for each stimulation condition over the four experimental runs of optokinetic stimulation: the mean intensity, the number and the mean duration of vection episodes. The mean intensity of vection sensation was calculated as the mean of the verbal report of vection experience (subjective scale, from 0 to 10). The mean duration of a vection period (self-motion perception) was calculated from the total time spent perceiving self-motion divided by the number of self-motion episodes reported.

#### Modeling of the electric field induced by tACS

The strength of the induced electrical field on the cortex surface was computed using biophysical modeling and checked for each participant. This measurement was used *a posteriori* to ensure that the electrical field on the cortical targets had a similar intensity and pattern to that simulated with the brain template for the tACS montage. To do so, participantś anatomic T1-weighted MRI was registered using a 3T scanner (Achieva 3.0T TX, Philips, Netherlands; T1TF2, TR = 25 ms, TE = 4 ms, voxel size = 0.95 mm 3 anisotropic). Head meshes of each participant were generated during the electrical field modeling with SimNIBS software version 4.0 (Nielsen et al., 2018; Thielscher et al., 2015). tACS modeling was done according to the quasi-static approximation as α-range is considered as a low frequency range.

### Statistics

We ensured that there were no violations of linearity concerning the mean intensity of vection (Kolmogorov-Smirnov test: p > .20) nor violation of sphericity (Mauchly test: p > .05). To assess the impact of stimulation condition on the mean intensity, repeated-measures analyses of variance (rmANOVA) with three stimulation conditions (α-tACS, θ-tACS, and sham) were performed. Post-hoc analyses were performed when necessary (Newman-Keuls test). Effect sizes are reported using partial eta-squared (*η_p_^2^*) for the main effects in rmANOVA, and Cohen’s d (*d*) for paired comparison. As neither the number of vection episodes nor the mean duration were normally distributed (Kolmogorov-Smirnov test: p < .20), Friedman test design was applied.

All statistical analyses were performed using Statistica software (v13.3, 1984–2017, TIBCO software INC, USA), with the significance level set at p < .05. Results and data distributions were displayed using Jasp software (JASP Team (2023)).

## Results

### Induced electrical field modeling and skin sensation

Among our participants, the simulated strength of the electrical field induced by the tACS montage reached an average of 0.59 V/m (range: 0.47 to 0.69 V/m). At the group level, the distribution of E-field strength was normal (Kolmogorov-Smirnov test: p > .20), with a standard deviation of 0.07 V/m.

Because the data distributions for the skin sensation score were not normal (Kolmogorov-Smirnov test: p < .20), Friedman test design was applied. The Friedman test was significant (χ^2^(2) = 17.71, p < .001). Participants reported slightly more unpleasant skin sensations when applying α-tACS (2.18 ± 0.99) or θ-tACS (1.98 ± 0.88) than for sham (1.23 ± 0.42). As revealed by Wilcoxon signed rank test analysis, there was no difference in skin sensation between α-tACS and θ-tACS (W = 270, p = .495), but both differed significantly from the sham condition (α-tACS vs. sham: W = 377, p < .001; θ-tACS vs. sham: W = 364, p < .001). However, these reported sensations remained quite low even under the α-tACS and θ-tACS conditions, as skin sensation was rated on a scale from 1, indicating no particular sensation, to 5, indicating a very unpleasant sensation to the point of wanting to stop the experiment.

### Behavioral data

Optokinetic stimulation induced a bistable perception in all the participants, with alternating periods during which they reported a sensation of self-motion and periods during which they perceived themselves as static. None of the twenty-two participants reported severe sickness symptoms as the rating of motion sickness did never exceed 6 on the scale ranging from 0 (“*no sickness*”) to 20 (“*severe sickness*”) as rated from the FMS.

#### Mean intensity of vection episodes

The rmANOVA revealed a significant effect of stimulation condition on the mean intensity of vection (F(2,42) = 7.393, p = .002, η_p_^2^ = .260). As can be seen in Fig. 3A, the mean intensity of vection was higher during α-tACS (6.28 ± 1) than during either θ-tACS (5.54 ± 2.13) or sham (5.01 ± 2.78). Neuman-Keuls *post-hoc* analysis confirmed that the mean intensity with α-tACS was significantly different from that with θ-tACS (p = .031, d = .511) and sham (p < .001, d = .776). In contrast, θ-tACS and sham did not differ from each other (p = .116). An additional correlation analysis indicated that there were strong linear and positive relationships between the three stimulation conditions (α-tACS vs. θ-tACS: r = .756; α-tACS vs. sham: r = .814; θ-tACS vs. sham: r = .823).

**Figure 3.**
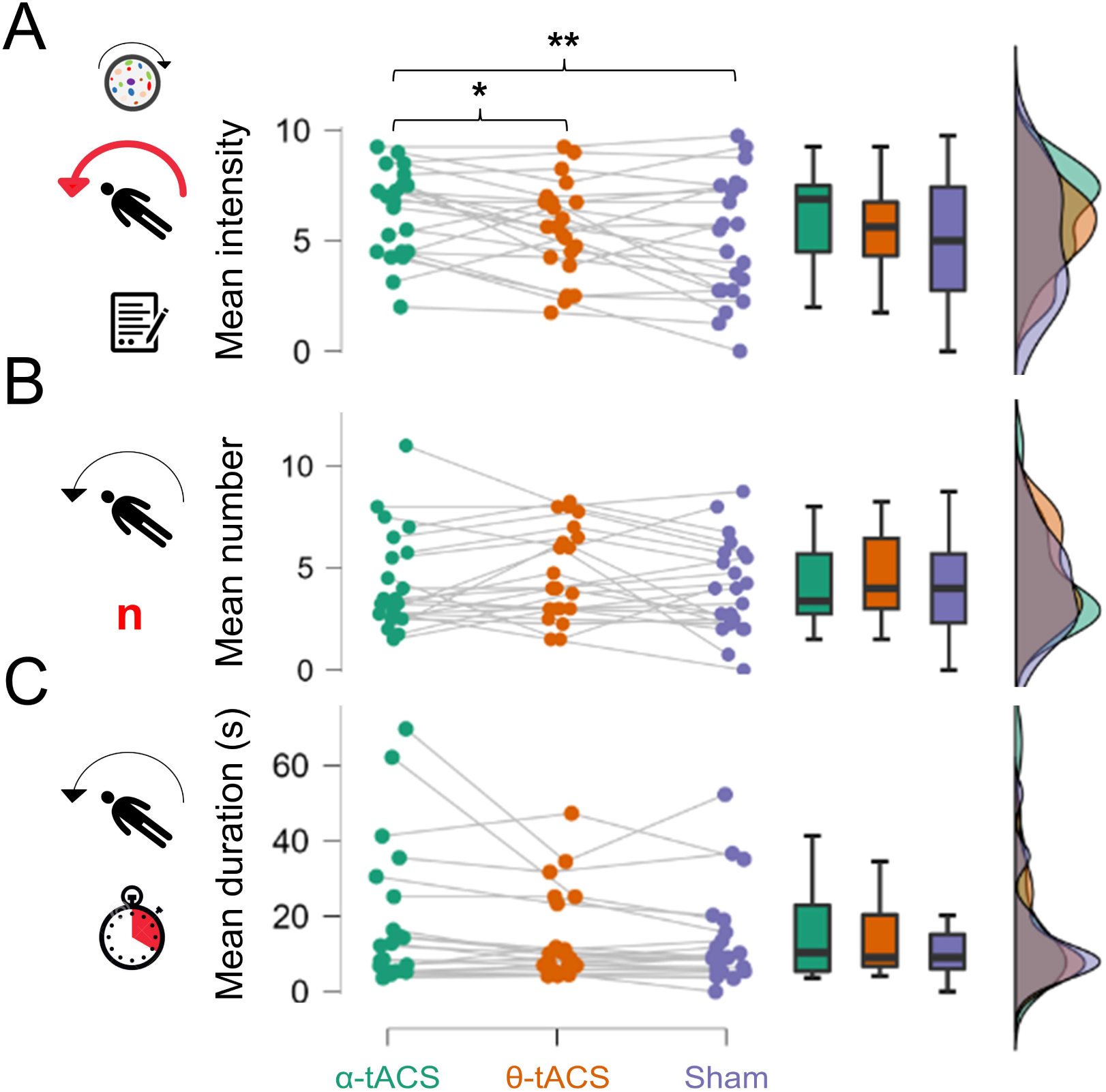
Modulation of vection quality by active and sham tACS. A: group-level distributions of vection intensity (subjective scale between 0 and 10) across the three stimulation conditions: α-tACS, θ-tACS and sham in green, orange and purple respectively. B: group-level distributions of the mean number of vection occurrences per 2 min run across the three stimulation conditions. C: group-level distributions of the mean duration of vection episodes (in s) across the three stimulation conditions. Significant post-hoc comparisons are indicated using stars (*: p < .05; **: p < .001).

#### Number and mean duration of vection episodes

The Friedman test showed that there was no effect of the stimulation condition on the number of vection occurrences (χ^2^(2) = 1.268, p = .530). The mean occurrence of vection for each run of two minutes was 4.30 (± 2.42) in the α-tACS stimulation condition, 4.72 (± 2.21) in the θ-tACS stimulation condition, and finally 4.05 (± 2.28) in the sham stimulation condition (Fig. 3B). This stability in the mean number of occurrences across stimulation conditions is accompanied by a fairly strong stability at the individual level, as shown in the correlational analysis (α-tACS vs. θ-tACS: r = .625; α-tACS vs. sham: r = .817; θ-tACS vs. sham: r = .623).

The results also showed that there was no effect of the stimulation condition on the mean duration of the vection period (χ^2^ (2) = 2.818, p = .244). Indeed, as can be seen in Fig. 3C, the mean duration of vection was fairly stable across stimulation condition, with a mean duration of 18.14 s (± 18.79 s) in the α-tACS condition, 14.00 s (± 11.84 s) in the θ-tACS condition and 13.55 s (± 12.62 s) in the sham condition (Fig. 3C). An additional correlation analysis indicated that there were strong linear and positive relationships between the three stimulation conditions (α-tACS vs. θ-tACS: r = .907; α-tACS vs. sham: r = .971; θ-tACS vs. sham: r = .909).

## Discussion

The present study is the first to have shown that visually induced self-motion illusions can be modulated by non-invasive brain stimulation. More precisely, our results show that the tACS-induced entrainment of the endogenous superior parietal α oscillations present during vection leads to an increase in the latter’s intensity. We hypothesize that this entrainment increases the strength of the inhibitory mechanisms operating during visuovestibular conflicts in the sensorimotor and vestibular functional networks. Lastly, our results suggest that the observed bistability phenomenon (as quantified by the number and duration of vection episodes) was not directly neuromodulated; this contrasted with the induced behavioral effect on the vection’s intensity.

We found that optokinetic stimulation induced bistable perception and thus produced alternating periods during which participants reported either a feeling of self-motion or of being static. Our main results suggest that neuromodulation of ongoing α oscillatory brain activity in the superior parietal cortex affects the quality of vection, when compared with control stimulation conditions that differed in terms of stimulation frequency (θ-tACS) or electrical intensity (sham). Overall, the illusion was more intense with α-tACS than with θ-tACS or sham stimulation (Fig. 3). Despite the lack of direct electrophysiological evidence, several points indicate that the observed behavioral effects were caused by the induced entrainment of endogenous α oscillations by tACS, Firstly, the results from the induced electrical field simulations confirmed that the average dose level (0.59 V/m) was nearly twice the minimum threshold of 0.3 V/m recently reported in the literature as inducing neurophysiological effects in awake, behaving mammals (Alekseichuk et al., 2022). Importantly, the low interindividual variability (0.07 V/m) showed that the participants had been stimulated equally. Secondly, dynamic systems theory predicts that neuronal modulation is most effective when the frequency of weak periodic perturbations is close to or at the resonance frequency of a brain network with intrinsic periodic dynamics (Ali et al., 2013; Liu et al., 2018). These dynamic properties are known to be present on the mesoscale in humans and differ from one lobe to another and from one Brodmann area to another; the dynamic signature of the parietal cortex is particularly sensitive to α-β oscillations (e.g. using TMS-EEG coupling (Darracq et al., 2018; Harquel et al., 2016; Rosanova et al., 2009)). In our study, alignment with the resonance frequency was guaranteed by tuning the α-tACS frequency to each participant’s individual frequency (IAF) (Krause et al., 2022). In a recent study of the awake ferret, Huang *et al*. (2021) reported that this phenomenon was particularly strong over the posterior parietal cortex. Individualized α-tACS was able to entrain spiking activity of individual cortical neurons during endogenous α oscillations, as shown by the spikes’ increased phase-locking to the stimulation oscillatory waveform (Huang et al., 2021).

The entrainment of α oscillations over the superior parietal cortex is therefore a plausible basis for the effects of the stimulation protocol used in this work. This entrainment echoes the increase in α activity within the same networks during visually induced vection, as reported recently by our group (Harquel et al., 2019) and by McAssey *et al*. (2020). The modulation of α activity (as measured in scalp EEG recordings) was thought to reflect the level of inhibition (Palva & Palva, 2007) within the sensorimotor and vestibular functional networks needed to reduce potential interference from conflicting vestibular inputs, which otherwise would reduce vection experiences (Harquel et al., 2019; Kleinschmidt et al., 2002). The tACS-induced entrainment of α oscillations would then trigger over-inhibition of these interferences and thus would ultimately increase the intensity of the illusion. Interestingly, we did not find evidence of neuromodulation of the number or duration of vection episodes during visual stimulation of the participants. This finding suggests that the entrainment of endogenous α oscillations was not strong enough to affect the transient decreases in the oscillatory activity (disinhibition) observed at vection onsets and offsets, which reflect the destabilization of percepts when entering into or exiting from vection (Harquel et al., 2019; McAssey et al., 2020). Again, the evidence suggests that tACS is effective in modulating ongoing neural oscillations. However, the ability of tACS to trigger robust exogenous modulation (particularly downmodulation) of oscillatory activity (as required during disinhibition phases) remains unclear (Huang et al., 2021; Liu et al., 2018).

As mentioned above, our results suggest that α-tACS modified the perceived intensity of the illusion, rather than modulating the bistable phenomenon. According to the literature, the amplitude of rotational displacements experienced using optokinetic stimulation in roll may be limited by the conflict between graviceptor signals and the visual effect (Dichgans & Brandt, 1978). However, our earlier research showed that some participants felt continuous self-rotation (Harquel et al., 2019). Coincidentally, these participants also displayed the greatest increase in α activity over the superior parietal cortex during visually induced vection. The effect of α-tACS on vection intensity observed in the present study maps well to the increased activation of the above-mentioned areas of the brain observed in previous studies of vection. The vection intensity measured in our study might be related to the feeling of presence (Held & Durlach, 1992; Sanchez-Vives & Slater, 2005), defined as the extent to which an artificial stimulus is interpreted as being real. The degree to which virtual reality, flight simulators and driving simulators provide participants with feelings of presence may have important implications for the application of these technologies (Palmisano et al., 2015; Stanney, 2014).

## Conclusion

To date, the evidence of the relationship between vection and parietal α oscillations has been correlative in nature. By using tACS to directly entrain endogenous α oscillations induced by vection over the parietal cortex, we observed a causal relationship between parietal α oscillations and vection quality for the first time. In the future, α-tACS might be of particular interest in virtual reality or flight/driving simulator applications and might enhance the feeling of presence associated with these technologies.

## Data availability statement

The data recorded in this study are available at https://osf.io/wepzc/ (DOI: 10.17605/OSF.IO/WEPZC).

## Author contribution

**Sylvain Harquel**: Conceptualization, Methodology, Formal analysis, Writing – original draft, Visualization, Supervision. **Corinne Cian**: Conceptualization, Methodology, Investigation, Formal analysis, Writing – original draft, Supervision. **Laurent Torlay**: Methodology, Investigation, Formal analysis, Writing – review & editing, Visualization. **Emilie Cousin**: Investigation. **Pierre-Alain Barraud**: Ressources. **Thierry Bougerol**: Ressources, Project administration. **Michel Guerraz**: Conceptualization, Methodology, Investigation, Writing – original draft, Supervision.

